# Deep brain stimulation creates information lesion through membrane depolarization

**DOI:** 10.1101/2022.03.26.485926

**Authors:** Eric Lowet, Krishnakanth Kondabolu, Samuel Zhou, Rebecca A. Mount, Cara Ravasio, Xue Han

## Abstract

Deep brain stimulation (DBS) is a promising neuromodulation therapy that alters neural activity via intracranial electrical stimulation. However, the neurophysiological mechanisms of DBS remain largely unknown, because of the difficulty of obtaining cellular resolution recordings without electrical interference. Here, we performed high-speed membrane voltage fluorescence imaging of individual hippocampal CA1 neurons during DBS in awake mice. We discovered that DBS, delivered at either 40Hz or 140Hz, reliably depolarizes somatic membrane potentials. Further, DBS enhanced spike rates and paced membrane voltage and spike timing at the stimulation frequency, though more prominent at 40Hz than 140Hz. To determine how DBS induced membrane voltage change impacts neuron’s ability to process inputs, we optogenetically evoked membrane depolarization. We found that neurons become unreliable in responding to optogenetic inputs during DBS, particularly during 140Hz DBS. These results demonstrate that DBS produces powerful membrane depolarization that interferes with neuron’s ability to process inputs, creating information lesion.

## Main

Deep brain stimulation (DBS) is effective in managing the neurological symptoms of Parkinson’s disease, essential tremor, and dystonia^1,2^. DBS directly stimulates brain tissue through chronically implanted electrodes and is considered a neural circuit specific therapy. Clinically, DBS is typically delivered at high frequencies of 130-200Hz, as lower frequency stimulations produce no consistent therapeutic effects^3–6^. The clinical success of DBS in managing movement disorders led to increasing effort in exploring the therapeutic benefits of DBS in many other neurological and psychiatric conditions. For example, high frequency 130Hz DBS of fornix, the major output of the hippocampus, is actively explored for Alzheimer’s disease^1,7^. However, the neurophysiological mechanisms underlying the therapeutic effects of high-frequency DBS remain unclear^7–12^.

DBS therapeutic outcomes and time courses are diverse and depend on the specific disease conditions targeted^13^. Since DBS effect in movement disorder is consistent with pharmacological lesion or surgical removal of the target brain tissue, DBS was first thought to inhibit local neural activity, likely via membrane depolarization induced action potential blockage or glia mediated adenosine release^14,15^. However, electrode recordings and biophysical modeling studies suggest that electrical pulses can directly excite axons leading to antidromic activation of neurons projecting to the stimulated area or orthodromic activation of downstream postsynaptic neurons. An alternative theory is that DBS entrains or paces neural activity^7^ which interferes with individual neuron’s responding to synaptic inputs and thereby creates information lesion^16^ that disrupts pathological network patterns. These theories have inspired recent exploration of stimulation waveform patterns and the development of closed-loop DBS devices that target pathological electrical field features in Parkinson’s disease and epilepsy^2^. While these theories are attractive, direct experimental testing of the neurophysiological effects of high-frequency DBS in the brain has been challenging due to electrical interference.

To probe the effect of DBS on individual neurons in real-time, we performed membrane voltage imaging of CA1 neurons, free of electrical interference, through chronically implanted optical imaging windows in awake mice^17^. AAV9-Syn-SomArchon-p2A-CoChR-Kv2.1 was infused through an infusion cannula coupled to the imaging window to express the genetically encoded voltage indicator SomArchon and the optogenetic actuator CoChR in the same CA1 neurons^17^. DBS was delivered via a stimulation electrode coupled to the imaging window, with the electrode tip positioned ~200μm below the imaging plane. A skull screw over the cerebellum was used as the ground (**Fig.1a&b**). We delivered DBS at 40Hz or 140Hz through the stimulation electrode, while performing SomArchon voltage imaging in mice freely navigating a spherical treadmill. DBS was delivered for 1 second, every 12 seconds, with each electrical pulse being 400μs width, bipolar, 10-60μA peak amplitude (38.3±11.4 μA, mean ± standard deviation, n=7 mice). Upon stimulation with either 40Hz or 140Hz DBS, we observed significant membrane voltage depolarization in individual neurons (**Fig. 1c&d**). To understand how DBS selectively impact action potentials versus subthreshold membrane voltage (Vm), we identified spikes and computed Vm by removing the identified spikes (detailed in Methods, **Extended data Fig.1**).

**Figure 1.**
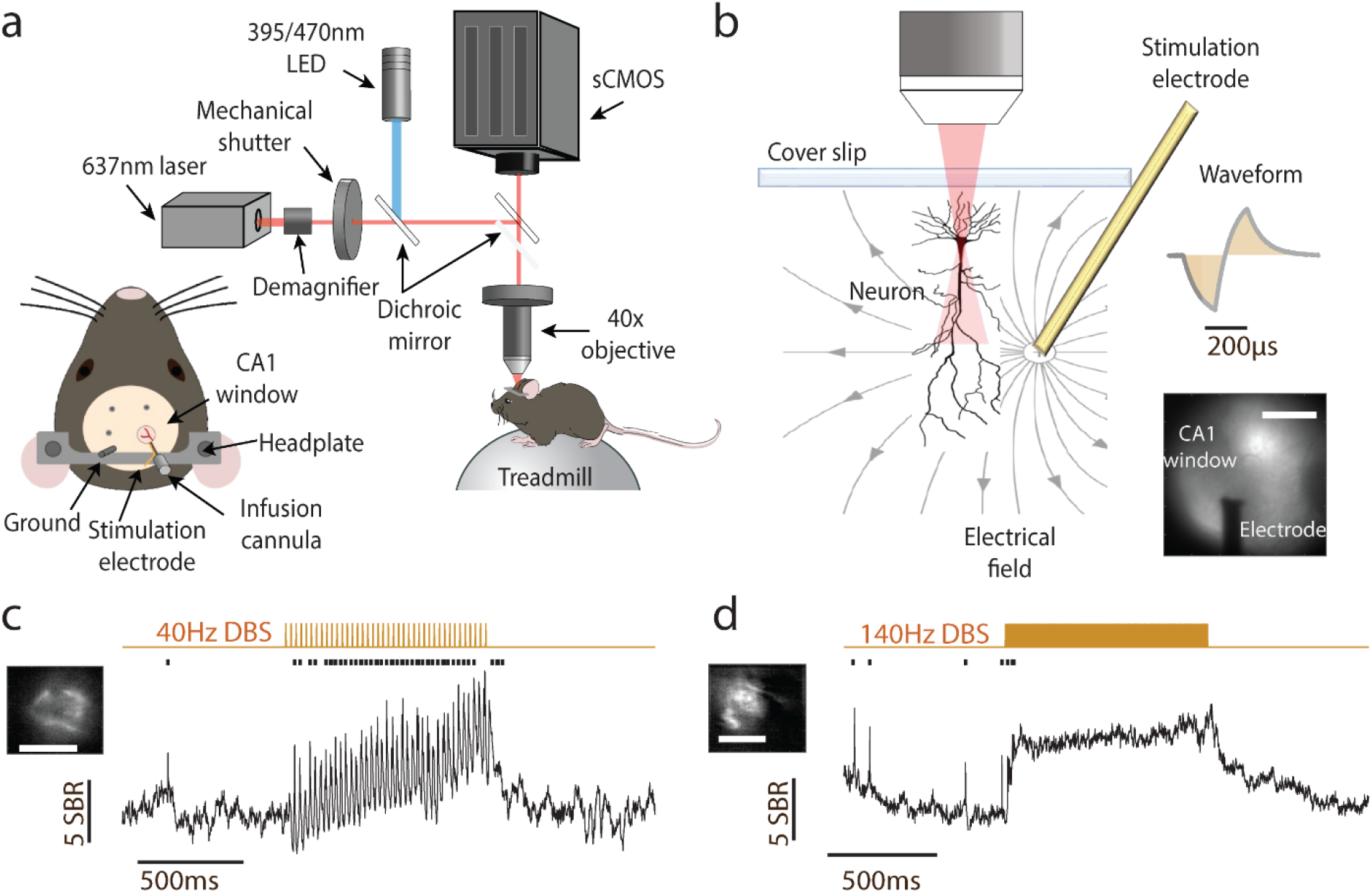
Single cell SomArchon fluorescence voltage imaging enables artefact-free neural recordings during DBS. **(a)**. Illustration of the experimental setup, the optical imaging window, and the animal preparation. **(b).** Schematic representation of a recorded CA1 neuron in the electric field generated by the electrode a few hundred microns away, and an example empirically measured electrical stimulation waveform. Lower right is an example imaging field showing GFP fluorescence from neurons expressing SomArchon-GFP and CoChR and the shadow of the nearby electrode. Scale bar 500μm. **(c).** Example SomArchon fluorescence before, during and after 40Hz DBS. Average SomArchon fluorescence of an example neuron is shown in the left upper image. Scale bar, 15μm. SomArchon trace is shown in black, detected spikes are marked by black ticks and electrical stimulation pulse patterns are in gold. **(d)**. Same as **c**, but for 140Hz DBS.

We found that Vm depolarization closely tracked the overall time course of the pulse trains, showing rapid depolarization at stimulation onset and rapid return to pre-stimulation baseline at stimulation offset (**Fig. 2a-c**). Interestingly, 40Hz DBS evoked Vm depolarization ramped gradually over the first couple hundred milliseconds following the onset, whereas 140Hz DBS evoked Vm depolarization rose to peak within a few milliseconds. Thus, we analyzed the initial transient response within 150ms of stimulation onset (transient period), and the sustained response during the remaining 150ms-1second period of each stimulation pulse train (sustained period). We found that 40Hz DBS led to significant Vm depolarization during both periods compared to the pre-stimulation baseline (paired t-test, df=20 neurons, transient: p=0.007; sustained: p=0.003). Similarly, 140Hz DBS led to prominent Vm depolarization during both periods (paired t-test, transient: p=1.69e^−4^, df=22; sustained: p=0.001, df=22). However, 140Hz DBS induced a significantly stronger depolarization than 40Hz DBS during the transient period, but not the sustained period (independent t-test, df=42, transient: p=0.011; sustained: p=0.45). Following stimulation offset, Vm quickly returned to the baseline level for both DBS conditions (paired t-test, 40Hz DBS: p=0.44, df=20; 140Hz DBS: p=0.91, df=22).

**Figure 2.**
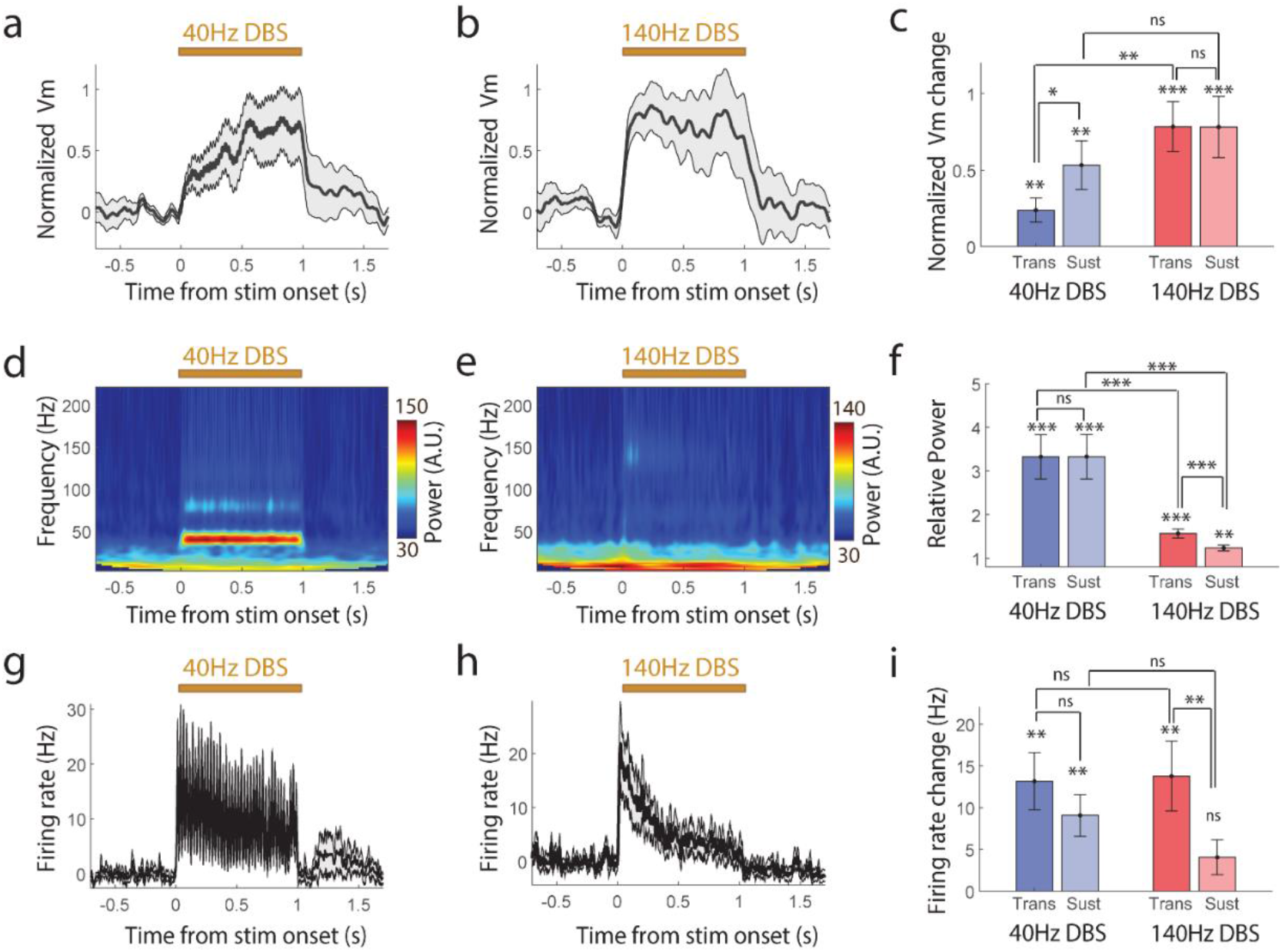
DBS induces powerful membrane depolarization modulations at both 40Hz and 140Hz. **(a-c).** Amplitude of the subthreshold membrane potential (Vm) and corresponding quantifications. **(a, b).** Population-averaged Vm during 40Hz DBS (**a**, n=21 neurons) and 140Hz DBS (**b**, n=26 neurons). **c.** Quantification of Vm change relative to the baseline during the transient (0-0.15sec) and sustained (0.15-1sec) periods of 40Hz and 140Hz DBS. **(d-f).** Time-frequency spectrum power of Vm and corresponding quantifications. **(d, e).** Population-averaged Vm power during 40Hz DBS (**d**, n=21) and 140Hz DBS (**e,** n=26). **(f).** Quantification of Vm 40Hz or 140Hz power change relative to the baseline during the transient and sustained periods of 40Hz and 140Hz DBS. **(g-i).** Spike rate and corresponding quantifications. **(g, h).** Population-averaged spike rate during 40Hz DBS (**g**, n=21) and 140Hz DBS (**h**, n=26). **i.** Quantification of spike rate change relative to the baseline during the transient and sustained periods of 40Hz and 140Hz DBS. ns=non-significant, **\***<0.05, **\*\***<0.01, and **\*\*\***<0.001.

Neurons are capable of following certain rhythmic synaptic inputs, a phenomenon known as entrainment, an important network communication mechanism. Entrainment by DBS has been proposed as a potential therapeutic mechanism, where DBS-mediated neural activity or entrainment interferes with neuron’s ability to process synaptic inputs leading to disruption of pathological network connectivity^16^. Thus, we next examined whether DBS could entrain or evoke precisely timed neuronal responses. We found that 40Hz DBS powerfully entrained Vm, leading to a prominent 40Hz component in Vm power spectrum (**Fig.2d**), which emerged at stimulation onset and sustained throughout the stimulation pulse train duration (paired t-test comparing to pre-stimulation baseline, transient: p=4.1e^−4^, df=19; sustained: p=5.17e^−4^, df=19, **Fig.2f**). 140Hz DBS also entrained Vm at 140Hz (paired t-test, transient: p=1.68e^−4^, df=21; sustained: p=0.006, df=21; **Fig.2e&f**), though the magnitude of 140Hz power induced by Vm entrainment at 140Hz was significantly weaker than the 40Hz power induced by Vm entrainment at 40Hz during both the transient and the sustained periods (independent t-test, transient: p=0.0013, df=19; sustained: p=2.14e^−4^, df=19; **Fig.2f**). Following stimulation offset, Vm power returned to the baseline level under both DBS conditions (paired t-test, 40Hz DBS: p=0.07, df=19; 140Hz DBS: p=0.16, df=21). Together, these results demonstrate that 40Hz DBS powerfully entrained Vm, where each electrical pulse within a pulse train reliably evoked Vm depolarization. 140Hz DBS had substantially weaker entrainment effect, but nonetheless evoked transmembrane voltage change that is time locked to stimulation pulses.

We then examined how DBS-induced Vm depolarization and entrainment influences spiking output. We found that 40Hz DBS led to a sustained increase in firing rate throughout the stimulation period (**Fig.2g**) (paired t-test compared to pre-stimulation baseline, transient: p=0.0011, df=20; sustained: p=0.0017, df=20). Following stimulation offset, spike rate quickly returned to the baseline level (paired t-test, p=0.45, df=20). Consistent with the prominent Vm entrainment effect, 40Hz DBS also entrained spike timing, quantified by spike phase locking value (PLV) relative to Vm oscillations at 40Hz (paired t-test, p=5.77e^−8^, df=19, **Extended data Fig.2**). Upon 140Hz DBS, we found a strong increase in firing rate during the transient period compared to the pre-stimulation baseline (paired t-test, p=0.003, df=25), followed by a tendency of weakly increased firing rate during the sustained period (**Fig.2h,** paired t-test, p=0.066, df=25). Additionally, we found a significant reduction in firing rate following 140Hz stimulation offset relative to the baseline (paired t-test, p=0.032, df=25). 140Hz DBS also led to a spike entrainment effect (paired t-test, p=2.1e^−4^, df=22), but significantly weaker than that observed with 40Hz DBS (independent t-test, p= 6.5e^− 6^, df=41, **Extended data Fig.2**). We found no significant difference in evoked firing rate change between 40Hz DBS and 140Hz DBS during both the transient and the sustained periods (independent t-test, transient: p=0.91, df=45; sustained: p=0.12, df=45). Together, these results demonstrate that slower 40Hz DBS substantially increased individual neuron’s spike rate and powerfully entrained both spike timing and Vm throughout the stimulation duration. In contrast, 140Hz DBS transiently increased spike rate and weakly entrained spike timing and Vm.

To understand how the observed DBS-induced neuronal depolarization influences the information transfer ability of individual neurons, we further characterized how DBS affects neuronal responses to optogenetic inputs. Specifically, we pulsed blue light at 8Hz to activate CoChR-mediated ion conductance while performing near-infrared SomArchon fluorescence voltage imaging of the same neurons and delivering DBS. For each 3-second-long trial, a neuron was optogenetically stimulated throughout the entire trial period, and 40Hz or 140Hz DBS was delivered for 1 second in the middle (**Fig.3a**). We found that blue light induced CoChR activation led to powerful membrane depolarization in individual CA1 neurons, and each light pulse evoked precisely timed spiking during the baseline period before DBS (**Fig.3b-d**). Both Vm and spikes followed rhythmic 8Hz CoChR activation, consistent with the general observation that CA1 neurons are easily entrained by theta frequency (~5-10Hz) inputs. During DBS, at 40Hz and 140Hz, we detected additional Vm depolarization on top of CoChR-evoked Vm depolarization, similar to that observed without CoChR activation (**Extended data Fig. 3a, Fig. 2d-f**). However, we failed to detect any additional spike rate increase in the presence of CoChR activation (**Extended data Fig. 3b**), in contrast to that observed without CoChR activation (**Fig. 2g-i**), suggesting that DBS modulation of spike rate depends on the activation state of the neuron.

**Figure 3.**
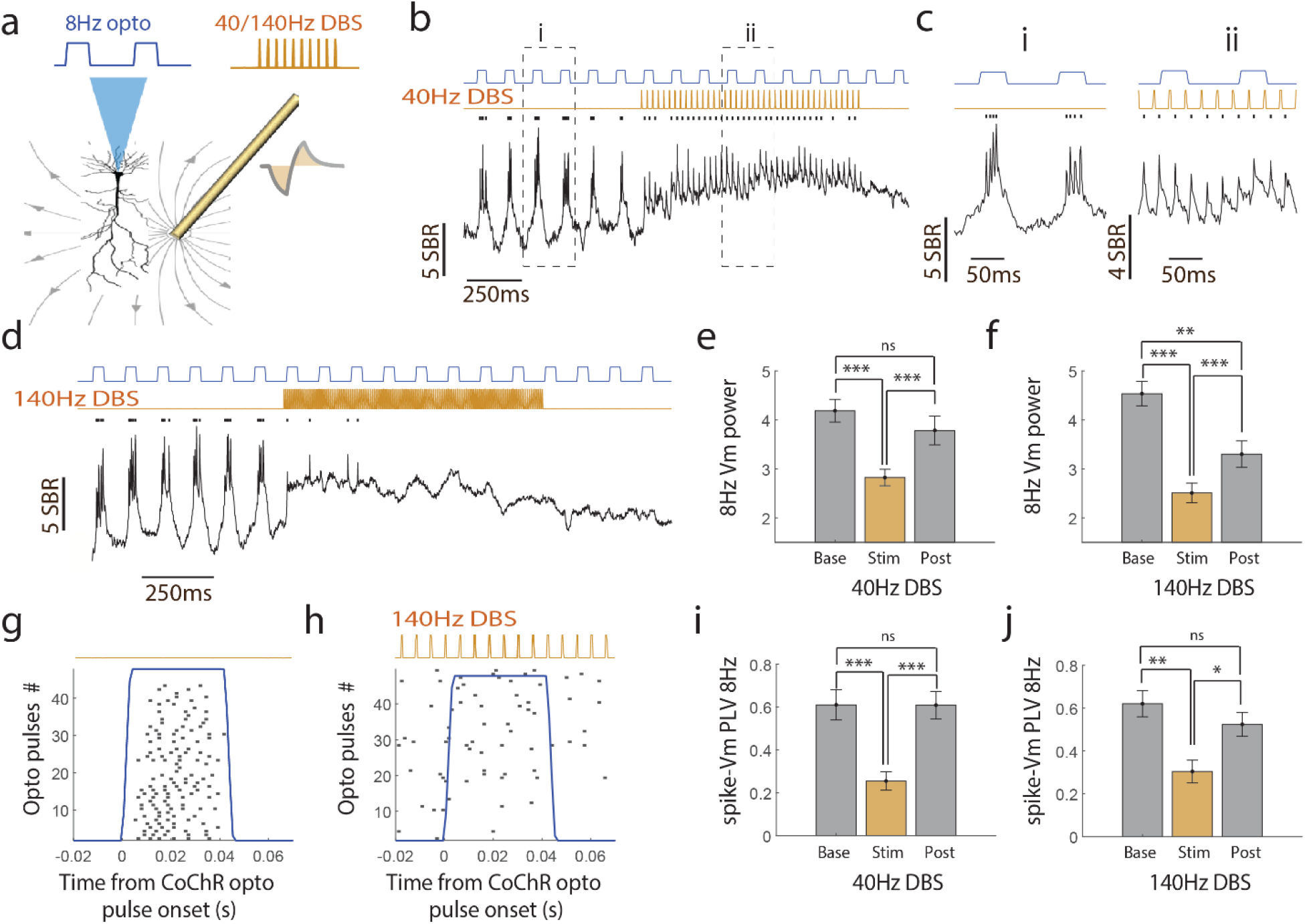
DBS reduces neuron’s ability to respond to optogenetically evoked depolarizing inputs. **(a)**. Illustration of simultaneous CoChR-evoked membrane depolarization and SomArchon voltage imaging during DBS. (**b, c).** An example CA1 neuron’s SomArchon fluorescence trace (black) and spikes (black ticks) during 8Hz CoChR activation (blue line) and 40Hz DBS (gold line). 8Hz CoChR activation occurred throughout the 3-seocnd trial, whereas 40Hz DBS occurred for 1 second in the middle of each trial. Zoom-in view (**c**) of the periods indicated by the dashed lines in (**b**), during the baseline (**i**) and the DBS period (**ii**). **(d)**. Same as in **b**, but for an example recording during 140Hz DBS. **(e)**. Quantification of 8Hz CoChR-evoked Vm power change during baseline (Base), 40Hz DBS (Stim) and post-stimulation periods (Post). **(f)**. Same as in (**e**), but for 140Hz DBS. (**g, h)**. An example neuron’s spike histogram aligned to blue-light pulse onsets delivered to activate CoChR during baseline (**g**) and 140Hz DBS (**h**). **(i)**. Quantification of spike phase-locking value (PLV) to 8Hz Vm during baseline (Base), 40Hz DBS period (Stim) and post-stimulation period (Post). **(j)**. Same as in (**i**), but for 140Hz DBS. ns=non-significant, **\***<0.05, **\*\***<0.01, and **\*\*\***<0.001.

To assess DBS effect on individual neuron’s information coding ability, we then analyzed the reliability of CA1 neuron in following 8Hz optogenetic inputs. Since Vm was reliably paced by 8Hz optogenetic activation of CoChR, we first computed 8Hz Vm power before, during, and after DBS (**Fig.3e**). With 40Hz DBS, we found a significant reduction of 8Hz Vm power during DBS compared to the pre-stimulation baseline (paired t-test, p= 3.23e^−7^, df=19), which largely recovered to the baseline level after DBS (paired t-test, p=0.075, df=19). With 140Hz DBS (**Fig.3f**), 8Hz Vm power decreased not only during DBS (paired t-test, p=3.42e^−7^, df=20), but remained suppressed after DBS compared to the baseline (paired t-test, p=5.96e^−4^, df=20). The reduction of Vm 8Hz power during 140Hz DBS was significantly greater than during 40Hz DBS (independent t-test, p= 0.046, df=39), and but not after DBS (independent t-test, p=0.055, df=39). Thus, DBS reduced Vm responding to rhythmic 8Hz inputs that are otherwise powerful at entraining CA1 neurons. Consistent with a loss of Vm entrainment to optogenetically induced 8Hz membrane depolarization, we found that during both 40Hz and 140Hz DBS, CoChR largely failed to evoke precisely timed spikes (**Fig.3h**). Spike-Vm PLV significantly reduced during DBS compared to the pre-stimulation baseline (**Fig.3i, j,** independent t-test, 40Hz DBS: p=3.23e^−4^, df=32; 140Hz DBS: p= 0.0023, df=29), which quickly recovered after DBS to the baseline level for both DBS conditions (independent t-test, 40Hz DBS: p= 0.99, df=24; 140Hz DBS: p= 0.37, df=25).

Finally, to understand how the loss of neuronal responding to inputs relates to DBS-induced membrane depolarization, we compared DBS effect on entrainment versus Vm depolarization. Across neurons analyzed, the average amplitude of DBS-induced Vm depolarization is correlated with the suppression of Vm entrainment by 8Hz optogenetic inputs (**Fig.4**, linear regression slope, r^2^=0.41, p<1e^−20^, n=40). Similarly, the amplitude of DBS-induced Vm depolarization is also correlated with the reduction of spike entrainment by optogenetic inputs (linear regression slope, r^2^=0.18, p=0.029, n=27). These results confirm that DBS-induced membrane depolarization is associated with suppressed responding of individual neurons to inputs.

**Figure 4.**
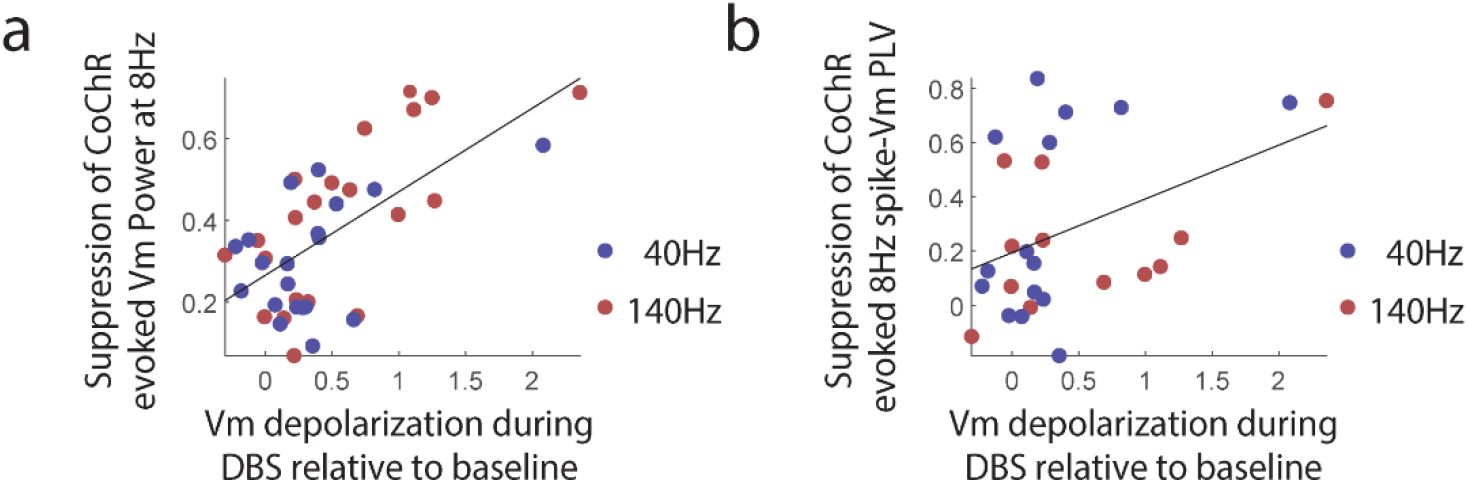
DBS induced membrane depolarization predicts suppression of optogenetic inputs. **a**. The reduction of optogenetically induced 8Hz Vm power is shown as a function of Vm depolarization amplitude during 40Hz (blue dots) and 140Hz (red dots) DBS stimulation. Each dot represents a neuron (n=40 neurons total). Neurons with stronger DBS-evoked Vm depolarization exhibits greater suppression of the 8Hz optogenetically induced Vm power. **b**. The reduction of optogenetically induced 8Hz spike-Vm phase locking value (PLV) is shown as a function of Vm depolarization amplitude during 40Hz (blue dots) and 140Hz (red dots) DBS stimulation (n=27 total neurons). Neurons with less than 5 spikes were excluded from this analysis.

In summary, we demonstrate that DBS powerfully depolarizes membrane voltage, with higher-frequency 140Hz DBS producing a more pronounced Vm depolarization than lower-frequency 40Hz DBS. Further, DBS increased spike rates and entrained spike timing throughout the stimulation period, particularly during the 40Hz DBS. Thus, our results do not support an overall neural silencing hypothesis, as we found continued spiking throughout the stimulation period. However, DBS, at both 40Hz and 140Hz, did suppress spike and Vm responding to optogenetic-evoked membrane depolarization inputs, and the magnitude of this suppression is correlated with the strength of DBS-induced Vm depolarization. Thus, DBS attenuates neuronal responses to inputs while depolarizing Vm and entraining spike output, which is in agreement with the information lesion hypothesis^16^. The substantial information blockade created by high frequency DBS, such as the 140Hz used here, could suppress pathological network synchronization and thus contribute to the network mechanisms of therapeutic DBS in Parkinsonian ^6,18,19^ or epilepsy patients^20^.

Different neurons with distinct biophysical properties will likely exhibit distinct entrainment effects to DBS of various frequencies. However, at higher frequencies, such as >130Hz, as typically used in the clinic, membrane ionic conductance time constants likely cannot support strong sustained entrainment. Thus, it is highly plausible that higher frequency >130Hz stimulation can reliably produce strong depolarization to create information lesion without strong entrainment, which might explain why high-frequency DBS is consistently effective across individual patients. The prominent entrainment properties of lower-frequency DBS may be beneficial in some conditions, such as the reported memory enhancement effect during 40-50Hz hippocampal DBS^20^. Future voltage imaging analysis of electrical stimulation effects in individual neurons across a wide range of brain regions will provide additional insights on how DBS affects neuronal responses across neural circuits and pathological conditions. Finally, future multi-neuron voltage imaging analysis could elucidate the network consequences of the DBS created information blockade effect observed here.

## Methods

### Animal preparation

All animal experiments were performed in accordance with the National Institute of Health Guide for Laboratory Animals and approved by the Boston University Institutional Animal Care and Use and Biosafety Committees. 7 female C57BL/6 mice (Charles River Laboratories, Inc.), 8-16 weeks at the start of the study, were used for all experiments. Mouse preparation was as described previously ^17,21,22^. Custom recording apparatus consists of an imaging window coupled with an infusion cannula (26G, PlasticsOne Inc., C135GS-4/SPC) and a stainless steel electrode for electrical stimulation (Diameter: 130μm, PlasticsOne Inc., 005SW-30S, 7N003736501F), using super glue (Henkel Corp., Loctite 414 and Loctite 713). The imaging window consists of a stainless steel cannula (OD: 3.17mm, ID: 2.36mm, 1.75mm height, AmazonSupply, B004TUE45E), with a circular coverslip (#0, OD: 3mm, Deckgläser Cover Glasses, Warner Instruments Inc., 64-0726 (CS-3R-0)) adhered to the bottom using a UV curable glue (Norland Products Inc., Norland Optical Adhesive 60, P/N 6001). The electrode tip protruded from the bottom of the imaging window by about 200μm, whereas the infusion cannula was leveled with the base of the imaging window.

Recording apparatus was surgically implanted under 1-3% isoflurane anesthesia, with sustained buprenorphine administered preoperatively to provide continued analgesia for 72 hours (buprenorphine hydrochloride, 0.03 mg/kg, i.m.; Reckitt Benckiser Healthcare). A craniotomy of ~3mm in diameter was made over the right dorsal CA1 (AP: −2mm, ML: +1.8mm). A small notch was made on the posterior edge of the craniotomy to accommodate the infusion cannula and the stimulation electrode. The overlying cortex was gently aspirated using the corpus callosum as a landmark, and the corpus callosum was carefully thinned to better expose the dorsal CA1. The imaging window was then positioned in the craniotomy, and Kwik-sil adhesive (World Precision Instruments LLC, KWIK-SIL) was applied around the edges of the imaging window to hold it in place. A small ground pin was inserted into the posterior part of the brain through the skull near the lambda suture, which was used as the ground for electrical stimulation. Three small screws (J.I. Morris Co., F000CE094) were screwed into the skull, and dental cement was then gently applied to affix the imaging window, the ground pin, and an aluminum headbar posterior to the imaging window. See **Fig.1a** for a diagram of recording apparatus placement.

AAV virus was infused via an infusion cannula (33G, PlasticsOne Inc., C315IS-4/SPC) connected to a microinfusion pump (World Precision Instruments LLC, UltraMicroPump3–4), through the implanted infusion cannula. Infusion cannula terminated about 200um below the imaging window. 1000nL of either AAV9-Syn-SomArchon-BFP-p2A-CoChR (titer: 1.53e^13^ genome copies (GC)/ml, Vigene Biosciences, Inc) or AAV9-Syn-SomArchon-GFP-p2A-CoChR (titer: 5.9e^12^ GC/ml, UNC vector core) was infused at a rate of 100nL/min, and the infusion cannula was left in place for another 10 minutes at the end of the infusion to facilitate AAV spread.

### Electrical stimulation

Electrical stimulation was delivered though an isolated pulse stimulator (Model 2100, A-M SYSTEMS). Stimulation consisted of 400μs bipolar pulses (negative phase= 200μs, positive phase= 200μs). We used either 40Hz or 140Hz pulse frequency. The peak amplitude per pulse ranged from 10-60μA (corresponding to 10-64μC/cm^2^ charge density per stimulation phase^23^) with a mean peak current of 38.3 μA and standard deviation of 11.4 μA across 7 mice. Stimulation pulse waveforms were calibrated with a function generator (TDS2022B, Tektronix), and one example waveform was shown in Figure 1A. Electrical stimulation sequences were externally triggered by TTL pulses generated by MATLAB (Mathworks Inc.) through a NI DAQ board (USB-6259, National instruments). TTL pulses were recorded at 10kHz sampling rate using the Open Ephys platform (http://open-ephys.org).

### SomArchon voltage imaging

Habituated mice were head-fixed on an air-pressured spherical Styrofoam ball and free to run. Animals were recorded 3-4 weeks after surgery. SomArchon imaging was acquired via a customized widefield fluorescence microscope equipped with a Hamamatsu ORCA Fusion Digital sCMOS camera (Hamamatsu Photonics K.K., C14440-20UP) and a 40x NA0.8 water immersion objective (Nikon, CFI APO NIR). A 140mW fiber-coupled 637 nm laser (Coherent Obis 637-140X) was coupled to a reverse 2x beam expander (ThorLabs Inc., GBE02-E) to obtain a small illumination area of ~30-40 μm in diameter to minimize background fluorescence. A mechanical shutter (Newport corp., model 76995) was positioned in the laser path to control the timing of illumination via a NI DAQ board (USB-6259, National instruments). The laser beam was coupled through a 620/60nm excitation filter (Chroma technology corp.) and a 650nm dichroic mirror (Chroma technology corp.), and SomArchon near-infrared emission was filtered with a 706/95nm filter (Chroma technology corp.). The fluorescence of GFP or BFP fused to SomArchon was used to localize SomArchon expressing cells during each recording. GFP was visualized with a 470 nm LED (ThorLabs Inc., M470L3), an 470/25 nm excitation filter, a 495 nm dichroic mirror and a 525/50nm emission filter. BFP was visualized with a 395nm LED (ThorLabs Inc., M395L4), a 390/18nm excitation filter, a 416nm dichroic mirror and a 460/60nm emission filter. SomArchon fluorescence was acquired at ~828 Hz (16 bits, 2×2 binning) using HCImage Live (Hamamatsu Photonics). HC Image Live data were stored as DCAM image files (DCIMG) and analyzed offline with MATLAB (Mathworks Inc.).

### Optogenetics

To excite CoChR, we used a blue 470 nm LED (ThorLabs Inc., M470L3) coupled to the widefield imaging setup with a 40x objective. 470 nm LED was controlled by a T-Cube LED driver (ThorLabs Inc., LEDD18, low gain) that was modulated using MATLAB (Mathworks Inc.) via NI DAQ board (USB-6259, National instruments). A neutral density filter (ThorLabs Inc., ND13A, optical density 1.3) was used to reduce the LED illumination density. We used a LED intensity range of 0.01-0.1 mW/mm^2^ depending on neurons’ response. Neurons were optogenetically activated by pulsed blue light at 8Hz (pulse width=42ms).

### SomArchon fluorescence images pre-processing and neuron identification

All offline analyses were performed with MATLAB (2019b&2020a, Mathworks Inc.). SomArchon fluorescence images were first motion corrected using a pairwise rigid motion correction algorithm as described previously ^24^. In short, the displacement of each image was computed by identifying the max cross-correlation coefficient between each image and the reference image. Our recordings consisted of multiple multi-second trials. We therefore concatenated all trials into a multi-trial image data matrix, and then we applied the motion correction algorithm. The motion-corrected image data were then used for subsequent manual neuron identification using the drawPolygon function (MATLAB). SomArchon fluorescence trace was extracted by averaging all the pixels within the identified neuron, and then detrended to correct for photobleaching using the function detrend (MATLAB). For detrending we only considered time points before or after DBS stimulation due to the large subthreshold modulations induced by electrical stimulation. The stimulation period was interpolated by averaging the 10 frames before and after the stimulation period making the detrending slope sensitive only to the baseline and post-stimulation fluorescence values.

### Spike identification and subthreshold Vm trace calculation

Spike detection was performed similar to that described previously in Xiao et al^25^. To identify spikes, SomArchon fluorescence trace were first high-pass filtered (>120Hz), and then spikes were detected as having fluorescence deflections greater than 4 standard deviations of the baseline fluctuations estimated from a spike-removed version of the high-pass filtered trace. To remove spikes, we first computed a smoothed trace by averaging the fluorescence trace using a moving window of ±100 frames. We then removed spike contributions from the SomArchon trace by replacing the fluorescence values above the smoothed trace with the corresponding smoothed trace values resulting in a spike-removed trace. To obtain the final subthreshold membrane voltage (Vm) traces, we removed three data points centered at the peak of each detected spike from non-filtered SomArchon trace, and interpolated the missing data points with the surrounding data points. To calculate spike signal-to-baseline-fluctuation ratio (SBR), we first obtained the spike amplitude by calculating the difference between the peak spike fluorescence and the lowest fluorescence value within three data points prior to the spike. We then divided the spike amplitudes by the standard deviation of the Vm across the entire recording duration.

### Spectral decomposition

Spectral decomposition of SomArchon Vm was performed with the FieldTrip Matlab toolbox^26^ (https://www.fieldtriptoolbox.org/), using wavelet morlet functions (5 cycles). The complex wavelets coefficients, from which we derived the phase and the amplitude, were used to compute spike-Vm phase locking values, and to estimate of the spectral power.

### Spike phase locking value (PLV) calculation

To obtain a measure of how consistent spikes occur relative to the phase of an oscillation we calculated the phase locking value^27^ (PLV), defined as:

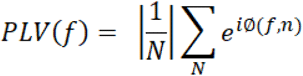

where the phase of a given frequency *f* was obtained from the complex wavelet spectrum of the subthreshold Vm or the optogenetic stimulation pulse train. We only included neurons that had more than 5 spikes.

## Acknowledgements

X.H. acknowledges funding from NIH (R01NS115797, R01NS109794, R01MH122971, and R01NS119483) and NSF (CBET-1848029 and 2002971-DIOS). E.L. acknowledges funding from Boston University Center for Systems Neuroscience. R.A.M. acknowledges NIH NRSA fellowship (F31MH123008). C.R. acknowledges funding from NSF GRFP. The funders had no role in study design, data collection and analysis, decision to publish, or preparation of the manuscript. We thank members of the Han Lab for technical support.

## Author Contributions

E.L., K.K, and S.Z. performed all imaging experiments. E.L. analyzed the data. R.A.M, E.L. and K.K. prepared the animals for the experiments. C.R. provided technical assistance. X.H. supervised the study. E.L. and X.H. wrote the manuscript. All authors edited the manuscript.

## Declaration of Interests

The authors declare no competing interests.

## Resource availability

**Lead Contact**

Further information and requests for code and data should be directed to the lead contact Xue Han.

## Materials availability

This study did not generate new unique reagents.

## Data and code availability

- Data are available from lead contact upon request.
- Codes used for data analysis is available on Github repository: https://github.com/HanLabBU.
- Any additional information required to reanalyze the data reported in this paper is available from the lead contact upon request.

## Extended data figures

**Extended data Fig.1.**
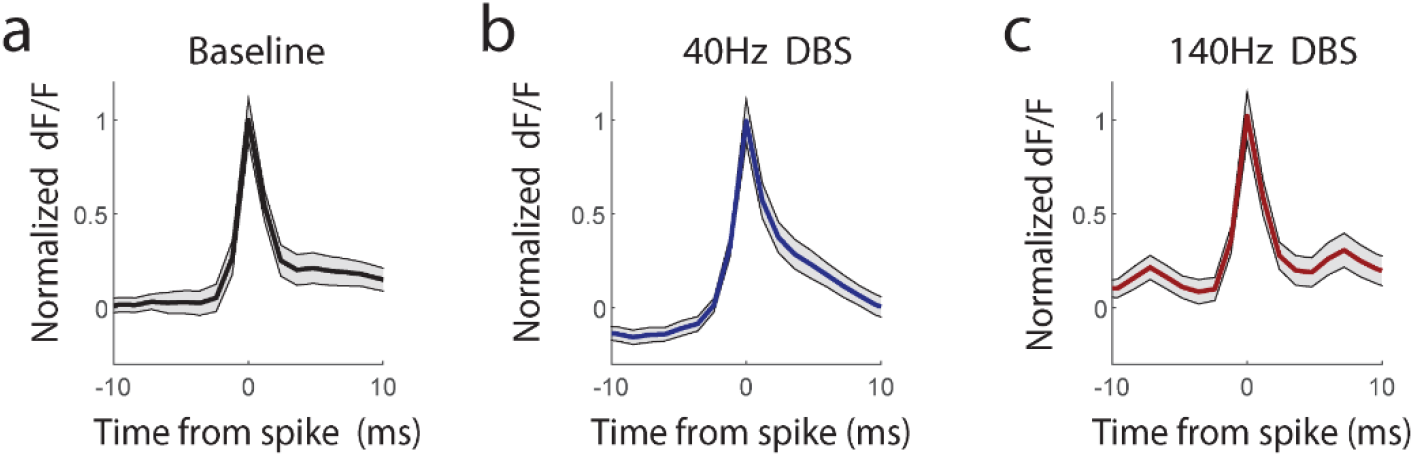
**a.** Normalized SomArchon fluorescence trace aligned to spike peak during the baseline period (the 1 second time window before DBS onset). SomArchon fluorescence for each neuron is obtained by first dividing by the averaged spike amplitude of the neuron and then subtracting the mean fluorescence during the −72ms to −12ms (50 frames) before spike peak. The center thick line is the mean SomArchon fluorescence across neurons, and the shading outlined with two thin black lines are ±standard error (n=21 neurons). **b**. Same as **a**, but for spikes occurring during 40Hz DBS. **c**. Same as **a**, but for spikes occurring during 140Hz DBS (n=26 neurons).

**Extended data Fig.2.**
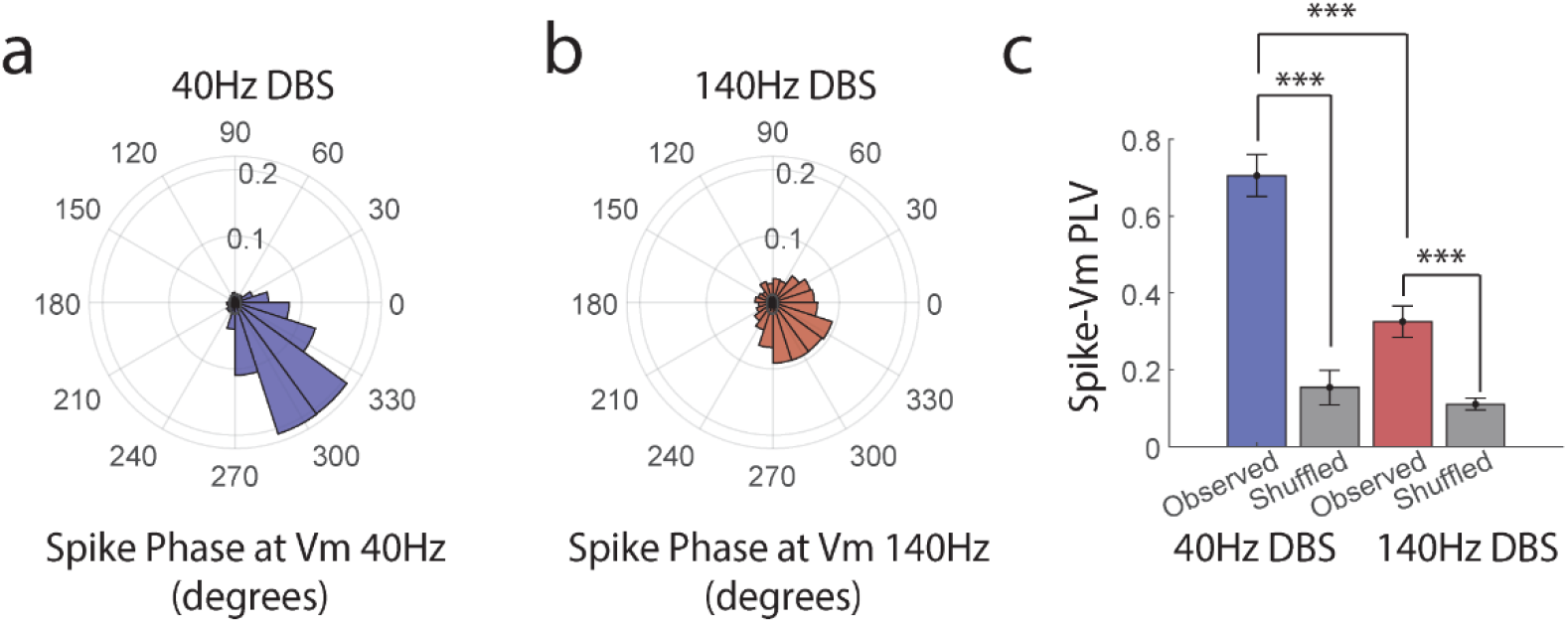
**a**. The population polar histogram of spike times relative to the phase of Vm filtered at 40Hz. Vm 40Hz oscillation phase was estimated from morlet wavelets (5 cycles). **b**. Same as **a**, but for 140Hz DBS. **c**. Quantification of the spike phase-locking values (PLV) relative to Vm during 40Hz or 140Hz DBS. To determine whether an observed PLV is significantly above a random distribution, we shuffled the spike times relative to the recorded Vm to obtain a PLV null distribution. ns=non-significant, **\***<0.05, **\*\***<0.01, and **\*\*\***<0.001.

**Extended data Fig 3.**
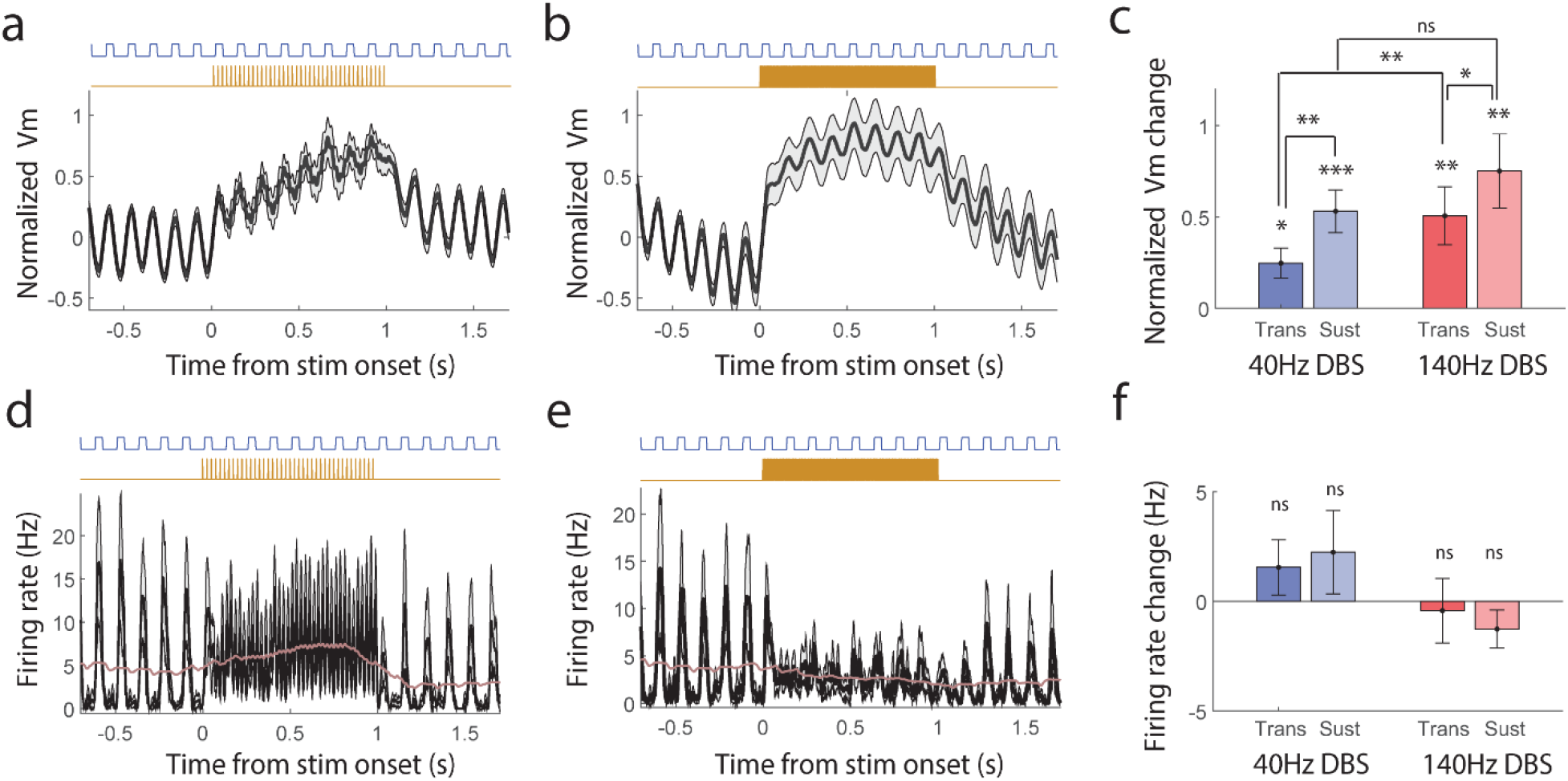
**a**. Population-averaged Vm during simultaneous 8Hz optogenetic activation (blue) and 40Hz DBS (gold). Vm is first normalized by the averaged spike amplitude in each neuron, and then averaged across neurons (n=16). **b**. Same as a, but with 140Hz DBS (n=17). **c**. Quantification of the transient (0-0.15sec) and the sustained (0.15-1sec) Vm depolarization induced by either 40Hz or 140Hz DBS, in the presence of optogenetic activation. **d**. Population-averaged firing rate during simultaneous 8Hz optogenetic activation (blue) and 40Hz DBS (brown, n=16 neurons). The purple line represents further smoothed firing rate (300ms rectangular smoothing). **e.** Same as **d**, but for population-averaged firing rate with 140Hz DBS (n=17). **f.** Quantification of the transient (0-0.15sec) and the sustained (0.15-1sec) firing rate changes relative to the baseline induced by either 40Hz or 140Hz DBS, in the presence of optogenetic activation. ns=non-significant, **\***<0.05, **\*\***<0.01, and **\*\*\***<0.001.

## Notes

### Competing Interest Statement

The authors have declared no competing interest.

